# SARS-COV-2 C.1.2 variant is highly mutated but may possess reduced affinity for ACE2 receptor

**DOI:** 10.1101/2021.10.16.464644

**Authors:** Xiang-Jiao Yang

**Affiliations:** Rosalind & Morris Goodman Cancer Institute, McGill University, Montreal, Quebec H3A 1A3, Canada; Department of Medicine, McGill University, Montreal, Quebec H3A 1A3, Canada; Department of Biochemistry, McGill University, Montreal, Quebec H3A 1A3, Canada; Department of Medicine, McGill University Health Center, Montreal, Quebec H4A 3J1, Canada

**Keywords:** COVID-19, coronavirus, virus entry, variant, B.1.1.7, B.1.351, P.1, N679K, NSP3, intra-host evolution, interhost evolution, nucleocapsid, C.1 variant

## Abstract

SARS-COV-2 evolution generates different variants and drives the pandemic. As the current main driver, delta variant bears little resemblance to the other three variants of concern (alpha, beta and gamma), raising the question what features the future variants of concern may possess. To address this important question, I searched through the GISAID database for potential clues. While investigating how beta variant has been evolving in South Africa, I noticed a small group of genomes mainly classified as C.1.2 variant, with one-year old boy identified in March 2021 being the index case. Over 80% patients are younger than 60. At the average, there are 46-47 mutations per genome, making this variant one of the most mutated lineages identified. A signature substitution is spike Y449H. Like beta and gamma variants, C.1.2 possesses E484K and N501Y. The genomes are heterogenous and encode different subvariants. Like alpha variant, one such subvariant encodes the spike substitution P681H at the furin cleavage site. In a related genome, this substitution is replaced by P681R, which is present in delta variant. In addition, similar to this variant of concern, three C.1.2 subvariants also encode T478K. Mechanistically, spike Y449 recognizes two key residues of the cell-entry receptor ACE2 and Y449H is known to impair the binding to ACE2 receptor, so C.1.2 variant may show reduced affinity for this receptor. If so, this variant needs other mutations to compensate for such deficiency. These results raise the question whether C.1.2 variant is as virulent as suggested by its unexpected high number of mutations.

## INTRODUCTION

Since the initial cases were identified in December 2019, coronavirus disease 2019 (COVID-19) has caused the global pandemic in a way totally unexpected to almost everyone in the world. The root driving force of this prolonged pandemic is continuous evolution of severe acute respiratory syndrome coronavirus 2 (SARS-CoV-2), with its first isolates sequenced and reported at the beginning of January 2020 [1]. The virus has evolved dynamically and yielded many variants, with some being the main drivers of the pandemic. Among them are four dominant strains, α (B.1.1.7) [2], β (B.1.351) [3], γ (P.1) [4] and δ (B.1.617.2) [5]. Without variants of concern and interest, the pandemic would have largely ended by the fall of 2020. Therefore, it is imperative to track how different variants evolve.

While trying to understand how β variant has been evolving in South Africa, I noticed a group of highly mutated genomes, many of which (but not all) have been designated as C.1.2 variant. Many of the C.1.2 genomes have been recently described by scientists from South Africa [6]. Like the four variants of concern, this new variant suddenly emerged from obscurity. Thus, an important question is whether this is a new candidate to be the next variant of concern. Here I present molecular and phylogenetic analyses of 130 C.1.2 genomes. Because one key substitution (Y449H) is known to reduce the affinity for the cell-entry receptor ACE2 [7], C.1.2 may need to lose this substitution or acquire additional mutations to compensate for the deficiency before it could become a new variant of concern. Related to the latter, a C.1.2-like genome, reported from South Africa on September 01, 2021, encodes the alarming substitution P681R, which is considered as a key substitution that makes δ variant so virulent.

## RESULTS AND DISCUSSION

### C.1.2 is one of the most mutated SARS-COV-2 variants

To understand how beta variant has been evolving in South Africa, I searched the GISAID sequence database (https://www.gisaid.org/) [8] for SARS-COV-2 genomes that were identified in the country after July 01, 2021 and encode the spike substitutions E484K and N501Y. Surprisingly, the search uncovered genomes not directly related to γ variant. Manual inspection of the listed mutations indicated that the genomes are related and encode the spike substitution Y449H. A majority of them received the Pango lineage designation C.1.2 (https://cov-lineages.org/) [9]. This designation was initiated on July 09, 2021 (https://github.com/cov-lineages/pango-designation/issues/139) and is thus relatively new. Because the parental C.1 lineage carries the signature NSP3 substitution Y428I, I searched through all SARS-COV-2 genomes from the GISAID database for those encoding both NSP3_Y428I and Spike_Y449H. The number of the uncovered genomes increased from 50 on August 10, 2021 to 62 on August 15, 2021 and 130 on September 02, 2021. As shown in Table S1, a majority of them has been designated as C.1.2, but nine genomes are classified as B.1.1.1, B.1.1.237, B.1.1.58, C.26 and C.37.1 [9]. Notably, the first case, identified in March 2021, received the designation B.1.1.1. The second case was identified on May 11, 2021 and is currently listed as the first case of the C.1.2 lineage (https://github.com/cov-lineages/pango-designation/issues/139), perhaps because the genome sequence of the March case was submitted to the GISAID database on August 26, 2021. Many of the C.1.2 genomes have been described in an important preprint released on August 24, 2021 [6]. As discussed below, the 130 genomes all belong to the same lineage. Also related is the new entry EPI_ISL_3859880, just reported from South Africa on September 01, 2021 and currently classified as B.1.1.373. It does not encode NSP3_T428I and was thus among the list of genomes encoding both NSP3_Y428I and Spike_Y449H.

Sex information is available for many of the patients associated with the 130 genomes encoding both NSP3_Y428I and Spike_Y449H, but no sex-specific association is evident. Among these 130 entries, 116 contain age information of the patients. Such information was downloaded from the database and used for generating Fig. 1A-B. Alarmingly, the patients are mainly under the age of 50. Of relevance, the patient associated with EPI_ISL_3859880 is 29 years old. Notably, the index case identified in March 2021 is 1-year old. CoVsurver, enabled on the GISAID website [8], was used to analyze the mutations in this case. As shown in Fig. 1C, the virus from the index case identified in March 2021 encodes many more mutations than the parental C.1 lineage. The mutation number distribution of the 130 genomes was generated via Coronapp, a web-based mutation annotation application [10,11]. As shown in Fig. 1D, each genome possesses an average of 46-47 mutations. This number is much higher than α [2], β [3] and γ (P.1) [12] variants of concern. In comparison, δ variant is much fast-evolving and has yielded different subvariants. The most mutated δ subvariants identified so far encode an average of 46-50 mutations per genome [13-16]. Thus, C.1.2 variant is one of the most mutated SARS-COV-2 lineages.

**Figure 1.**
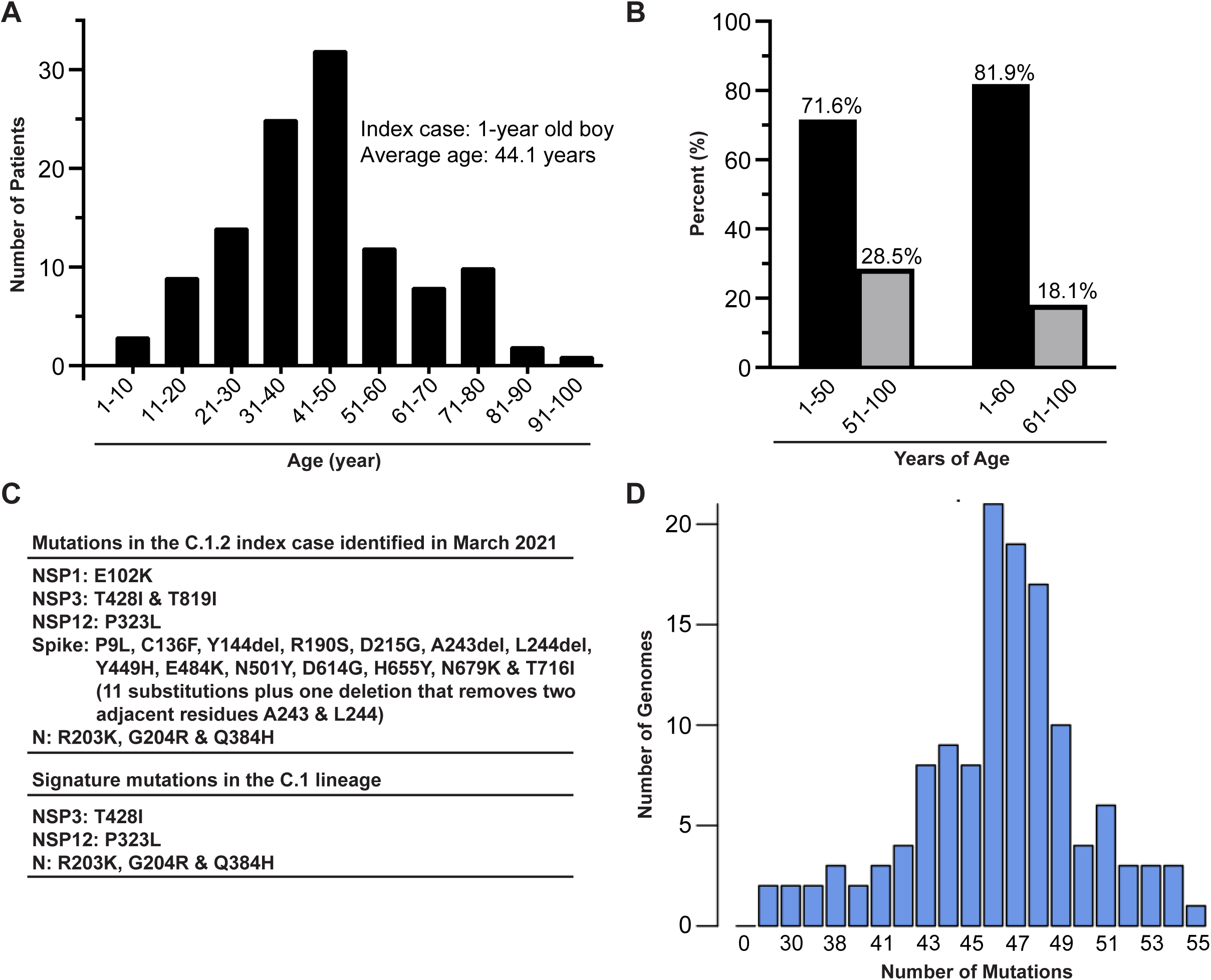
Age distribution of C.1.2 clinical cases and their average mutation load. (**A-B**) The NSP3 substitution Y428I and the spike substitution Y449H were used to search through over 3 million SARS-COV-2 genomes from the GISAID database on September 02, 2021. This search yielded 130 genomes, with 116 of them containing information on age and gender. Such information was downloaded from the database and used for generated panels (A) and (B). (**C**) CovSurver (https://www.gisaid.org/epiflu-applications/covsurver-mutations-app/) was used to analyze the mutations in the index case that was identified in South Africa in March 2021. (**D**) Mutation load of the 130 genomes. The distribution was generated via Coronapp (http://giorgilab.unibo.it/coronannotator/), a web-based mutation annotation application. The source is gratefully acknowledged in Table S1.

Its high mutation load raises the important question whether this variant is also highly virulent. To address this question, I quantified the monthly genomes sequenced in South Africa and deposited into the GISAID database (accessed on October 13, 2021). In June, July and August 2021, the C.1.2 genomes were 112, 85 and 31, respectively. During the same period, the δ genome number increased from 1,971 to 2,141 whereas the β genome number decreased from 532 to 8. In September 2021, there were 669 δ genomes, 18 C.1.2 genomes and only one β genome identified in South Africa. These results indicate that C.1.2 is more virulent than β variant but much weaker than δ variant. Therefore, C.1.2 variant may not as virulent as its high mutation load suggests. One question is whether there is any molecular basis for this.

### C.1.2 genomes are heterogeneous and form different groups

To determine the C.1.2 mutation profile, the 130 genomes from the NSP3_T428I and Spike_Y449H search were analyzed via Coronapp for mutations in all SARS-COV-2 genes. Shown in Fig. 2 are substitutions in NSP3, spike and nucleocapsid proteins. As expected, all genomes encode the NSP3 substitution T428I (Fig. 2A). A majority also encodes T819I of NSP3. In addition, there are two small groups, with one encoding P822S while the other encodes both T237I and H1274Y (Fig. 2A & Table S2). There are also silent mutations that affect the codons of F106, K1660 and S1882, which might affect mRNA stability and/or translational efficiency. As stated above, the entry EPI_ISL_3859880 does not encode T428I. But this genome does share the substitution T819I with many of the 130 genomes. In addition, it encodes P1228L. Notably, among these NSP3 substitutions, both P822S and P1228L are recurrent in two groups of delta subvariants [15], suggesting the potential importance. Due to the focus on the spike protein, other SARS-COV-2 proteins such as NSP3 have been ignored in many studies. However, NSP3 is a large multidomain protein with an important role in cytoplasmic double-membrane vesicles important for virus replication [17]. Moreover, this gene is one the most frequently mutated target. Thus, the NSP3 substitutions may be important in improving viral fitness.

**Figure 2.**
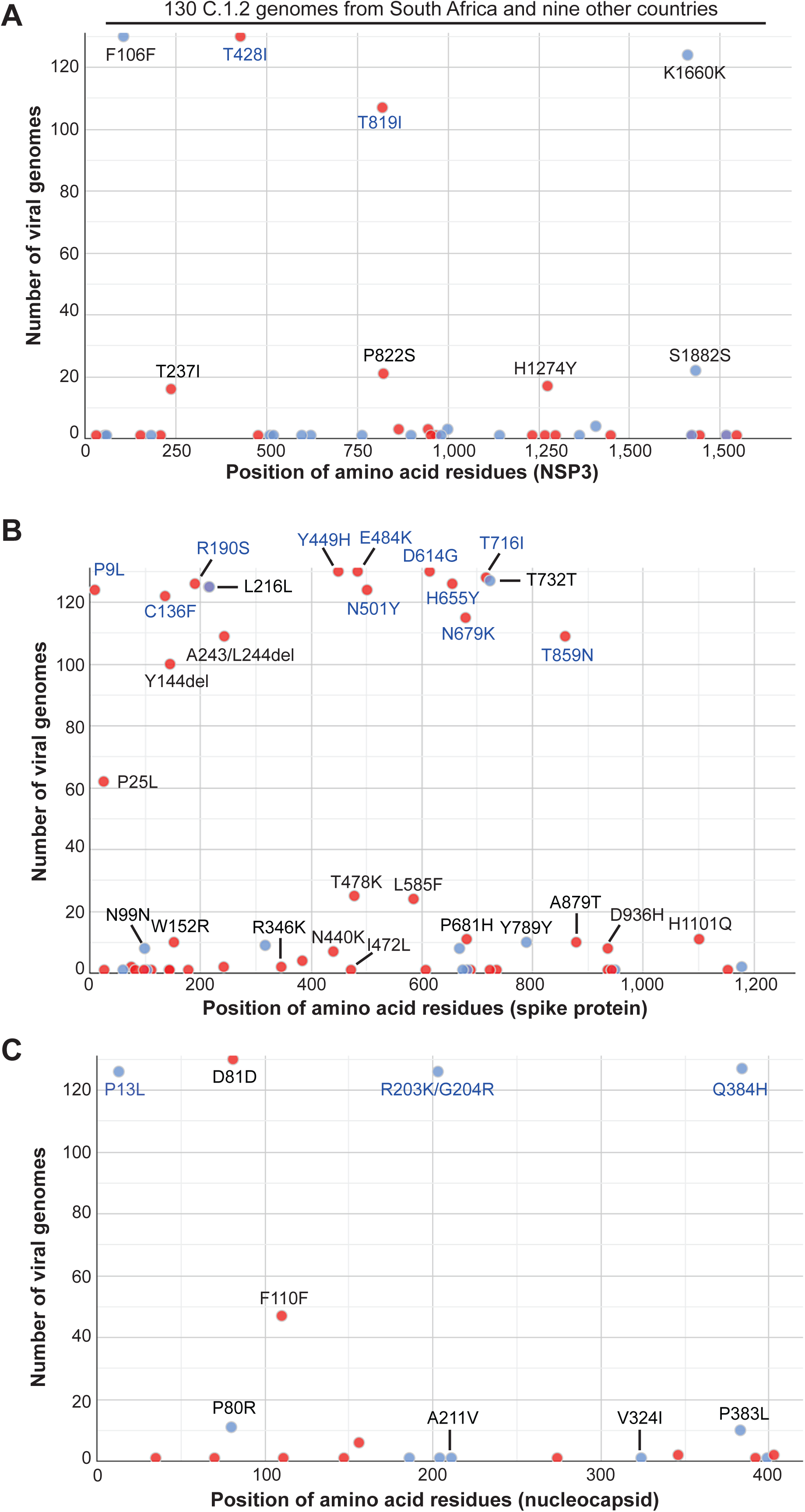
Mutation profile of 130 C.1.2 genomes from South Africa and nine other countries around the world. As in Fig. 1D, the genomes were downloaded from the GISAID database on September 02, 2021 for mutation profiling via Coronapp. Shown in (A), (B) and (C) are substitutions in NSP3, spike and nucleocapsid proteins, respectively.

As shown in Fig. 2B, almost all 130 genomes encode 9 spike substitutions: P9L, C136F, R190S, D215G, Y449H, E484K, N510Y, D616G and T716I. These substitutions are enriched in the N-terminal half of the protein (Fig. 1E). N679K is encoded by 115 genomes, while 109 genomes encode T859N and the deletion A243/L244. Another deletion, Y144, is present in 100 genomes. P25L is encoded in 62 genomes. Mutually exclusive, the substitutions T478K and L585F are present 25 and 24 genomes, respectively. Among those lacking N679K, 11 genomes encode the substitution P681H, with three associated with T478K and 7 linked to N440K in 7 genomes. Two in the former group contain the substitution R346K and the two deletions (Y144del and A243L244del, Fig. 1C). The latter group is also associated with D936H. The entry EPI_ISL_3859880 encodes the 9 signature spike substitutions but not P25L or T859N. Instead of N679K and P681H, it encodes the substitution P681R. These three substitutions are located within the furin recognition site (Fig. 1F). Thus, the 131 genomes form different groups and within each group, there is also a certain degree of variation. This mutational profile reiterates the heterogeneity of the spike protein encoded by the genomes. The heterogeneity is clearly much more evident than that in NSP3 (Fig. 2A).

Compared to the spike protein, the mutation spectrum of nucleocapsid protein is much less complex (Fig. 2C). Almost all genomes encode the substitutions P13L, R203K, G204R and Q384H. In addition, 11 and 10 genomes encode the substitutions P80R and P383L, respectively. They are mutually exclusive in two different groups. Like A211V, V324I is present in present in one genome. In addition to NSP3, spike and nucleocapsid proteins, substitutions are present in other viral proteins. But the variation is much less. Therefore, the gene for spike protein is the most variable among the 131 genomes, reiterating that this protein is the most frequent target SARS-COV-2 utilizes to improve its viral fitness. These results also suggests that C.1.2 variant has evolved and yielded different groups of subvariants.

### Phylogenetic analysis identifies different C.1.2 subvariants

Next, phylogenetic analysis was used to characterize these subvariants. For this, the software package RAxML-NG [18] was employed to analyze the 130 genomes. This analysis generated 20 maximum likelihood trees and one bestTree. Figtree was used to display these 21 trees for manual inspection and re-rooting. Presented in Fig. 3 is one re-rooted maximum likelihood tree with the initial three genomes from March and May 2021 settled at the two root branches. As evident from Fig. 3, the index case is part of a group that lacks the substitutions P25L, Y144del and T859N, whereas the two cases from May 2021 are part of a different group that possesses these alterations along with the substitution L585F. The T478K substitution is present in two small branches, with one encoding N679K and the other possessing P681H. The latter substitution is also present in a small branch that encodes N440K. EPI_ISL_3859880 encodes P681R, instead of N679K or P681H. Phylogenetic analysis placed this genome next to EPI_ISL_3799035 in the group encoding Y144del but lacking the P25L substitution (Figs 3 & S1). Because P681H and P681R are mutually exclusive with N679K, an interesting question is how P681H and P681R were introduced. The presented tree suggests that simultaneous gain of P681H/R and loss of N679K. While this could occur via recombination, it is also possible that the root of the P681H (or P681R) branch is much earlier than what suggested by the presented tree. Thus, there is still a certain degree of uncertainty about the phylogenetic relationship.

**Figure 3.**
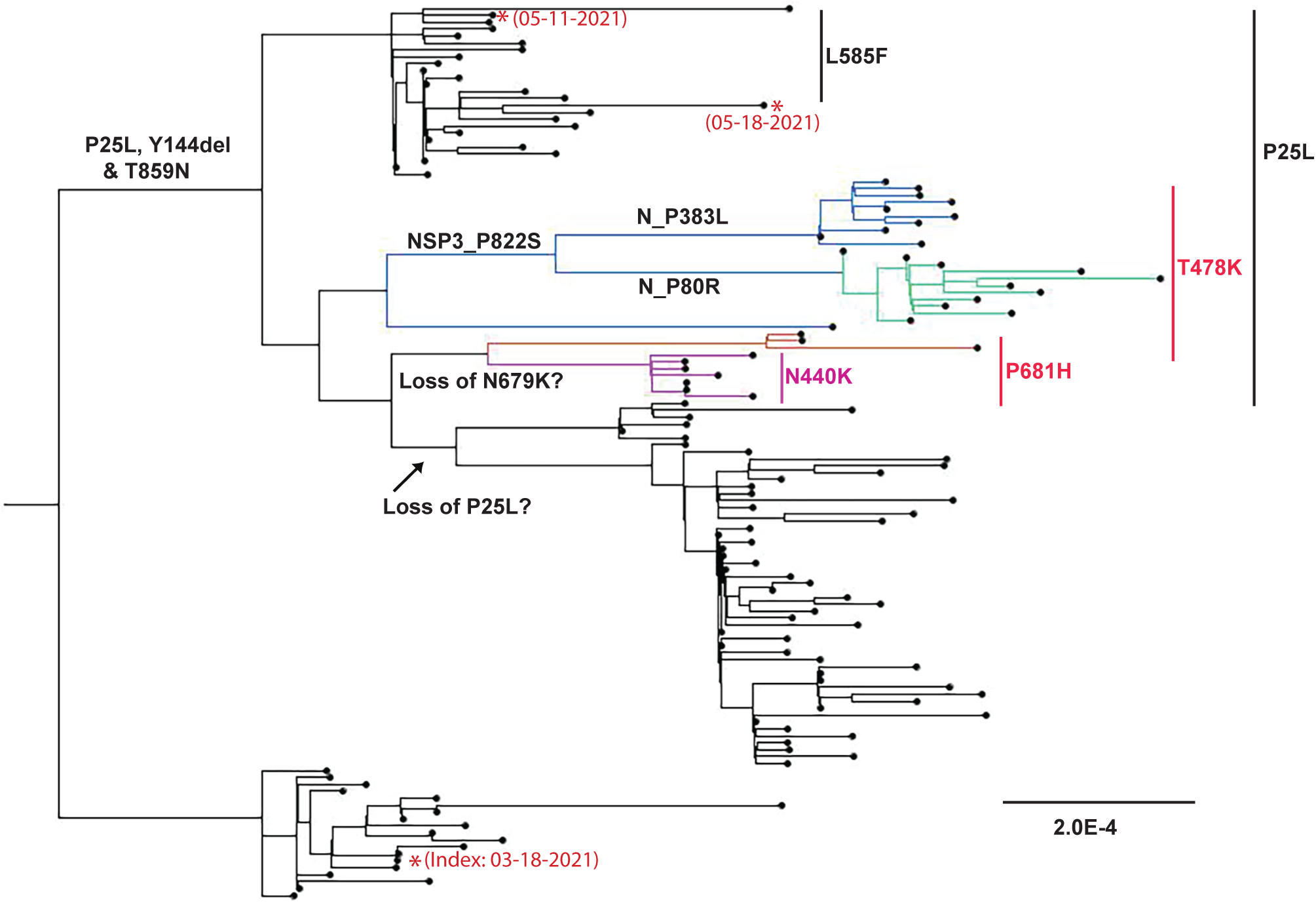
Phylogenetic analysis of the 130 C.1.2 genomes. The phylogenetic analysis package RAxML-NG was used to generate 20 maximum likelihood trees and one best tree. Figtree version 1.4.4. was used to display these 21 trees for manual inspection and tree re-rooting. Presented here is one re-rooted maximum likelihood tree with the initial index genomes from March and May 2021 at the two root branches. The strain names and GISAID accession numbers of the genomes are provided in Figure S1.

The presence of different subvariants begs the question about their potential trajectories in the new cases identified. According to the time when the cases corresponding to the available genomes were identified, the T478K branches are most frequently associated with the new genomes identified in August 2021, raising the possibility that these branches are the most active buds. This is also consistent with the fact that this substitution confers clear advantage to viral fitness. These branches encode either N679K or P681H. Compared to the former, P681H is more likely to improve furin cleavage [19]. In this regard, P681R encoded by the EPI_ISL_3859880 is more advantageous than P681H for the virus and thus deserves more attention.

### Functional impact of different spike and nucleocapsid substitutions

Two interesting questions are whether and how different mutations may exert functional impact. As shown in Fig. 1E, many spike substitutions are located within the N-terminal half that comprises the N-terminal domain and the receptor-binding domain. The substitution P9L is located with the signal peptide and may thus affect spike protein maturation and trafficking. P25L, C136F, Y144del, W152R, R190S, D215G are A243/L244del are located within the N-terminal domain. Except for C136F, similar substitutions have been reported for for alpha, beta and mamma variants. As shown for those these three variants, P25L, C136F, Y144del, W152R, R190S, D215G are A243/L244del are likely to confer immune evasion. As shown in Fig. 4A, C136 forms a disulfide bond with C15 (Fig. 4A), so the substitution C136F disrupts this disulfide bond and may thus affect the structure of C15, C136F and their neighboring regions. As these regions are also potential immune epitopes, C136F may also confer immune evasion. The signature substitution Y449H alters a key residue in the receptor-binding pocket [7,20,21]. As shown in Fig. 4B, Y449 is in proximity to D38 and Q42 of ACE2. Hydrogen bond formation of Y449 with D38 or Q42 of ACE2 contributes significantly to the receptor recognition by spike protein. Indeed, Y449H has been shown to reduce receptor binding substantially [7]. Thus, it is very likely, this substitution is disadvantageous for receptor recognition. However, Y449 is also recognized by antibodies from infected patients [22], so Y449H is advantageous in terms of immune evasion. This dilemma suggests that C.1.2 variant acquired this substitution under immune selection. It is noteworthy that the molecular studies of Y449H were carried out with the parental strain identified at the initial stage of the pandemic [7] and does not contain substitutions E484K, N50Y and D614G in C.1.2 and three variants of concern. Among the residues affected by these three substitutions, N501 is only 7.1 **Å** away from Y449 (Fig. 4C). Whether the impact of Y449H is different in variants that encode N501 remains to be investigated.

**Figure 4.**
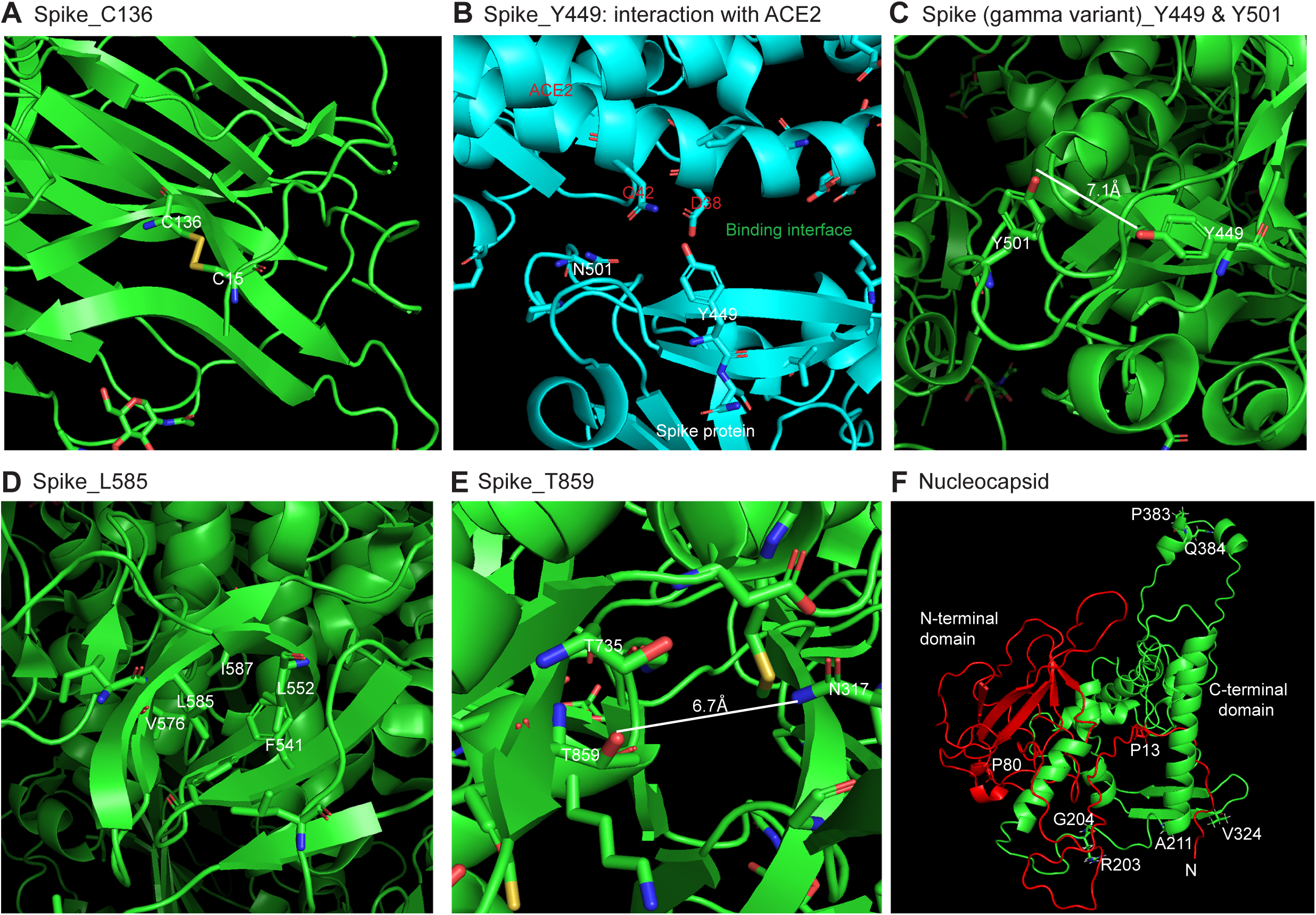
Potential functional impact of different spike and nucleocapsid substitutions. (**A**) C136 forms a disulfide bond with C15. The structures are PyMol presentation of the original spike protein structure 6XR8 from the PDB database. The alpha spike protein shows the same disulfide bond (PDB, 7KRQ). (**B**) Structural details showing spike Y449 and its proximity to D38 and Q42 of ACE2. The structure is PyMol presentation of the spike protein-ACE2 complex structure 6M17 from the PDB database. (**C**) Spike Y449 is 7.1 **Å** away from N501. The structure is PyMol presentation of the gamma spike protein structure 7M8K from the PDB database. (**D**) Structural details showing spike L585 and its hydrophobic proximity. Based on the gamma spike protein structure 7M8K from the PDB database. (**E**) Spike T859 is 6.7 **Å** away from N317. Based on the alpha spike protein structure 7KRQ from the PDB database. (**F**) Structure of nucleocapsid protein of SARS-COV-2. While R203 and G204 are in the serine/arginine (SR)-rich loop that connecting the N- and C-terminal domains, P383 and Q384 are within a bipartite nuclear localization signal sequence located at the C-terminal tail. The SR domain structure is hypothetical and the entire nucleocapsid structure is based on PyMol presentation of a structural model, QHD43423.pdb, built by Dr. Yang Zhang’s group at University of Chicago (https://zhanglab.ccmb.med.umich.edu/COVID-19/) from crystal structures of the N- and C-terminal domains of nucleocapsid protein (6M3M and 6YUN from the PDP database, respectively). N, N-terminal end.

Some subvariants encode T478K, also a signature substitution of delta variant that may confer some evolutionary advantage in terms of receptor expression and binding [7]. Moreover, one genome also encodes R346K, known to enhance receptor expression and binding [7]. Moreover, this and two other T478K-encoding subvariants carry P681H, which is expected to improve the furin recognition site (Fig. 1F). In addition to these three genomes, another small group encodes P681H (Fig. 3). Instead of T478K, these genomes encode N440K (Fig. 1F). In addition, two other groups of genomes encode T478K. Instead of P681H, N679K alters the furin recognition site. N679 and P681 are residues 7 and 5 N-terminal from the cleavage site, respectively. According to consensus sequence analysis [19], residue 5 is more important than residue 7, so N679K is expected to exert smaller effects than P681H. Because an arginine residue is much preferred than histidine and any other residues, P681R improves furin cleavage, as reported recently [23]. P681R may also contribute to membrane fusion by delta variant [24]. EPI_ISL_3859880 encodes P681R and thus deserves attention about its evolution further.

Shown in Fig. 4D are structural details of spike L585 and its neighboring residues. Many of them are hydrophobic, so L585F may improve such hydrophobic core. Spike T859 is 6.7 **Å** away from N317 (Fig. 4E), so T859N may confer interaction between N859 and N317. Other spike substitutions such as H655Y and T716I are recurrent in other major variants, so they may thus offer additional advantages. Thus, it is likely that except for Y449H, many spike substitutions are clearly advantageous in improving viral fitness.

Four substitutions, P13L, R203, G204 and Q384H, are present in all C.1.2 genomes. The latter three are signature substitutions of the C.1 lineage. R203, G204 are in the serine/arginine (SR)-rich loop connecting the N- and C-terminal RNA-binding domains, whereas Q384 is within a bipartite nuclear localization signal sequence located at the C-terminal tail (Fig. 4F). P13 is located at the N-terminal tail, which is a known immune epitope, so P13L may confer immune evasion. Two small groups of genomes encode P80R or P383L. P80 is located in the N-terminal domain and is a signature substitution of gamma variant, while P383 is within a bipartite nuclear localization signal sequence located at the C-terminal tail. One genome encodes A211V is C-terminal from this SR loop.

### Emergence of C1.2 variant via accelerated intrahost evolution

Schematic representation in Fig. 5A summarizes how different C.1.2 variant may evolve and gain additional substitutions. The 130 C.1.2 genomes belong to different subvariants, with one large group possessing N679K and a small group containing P681H. These two substitutions are mutually exclusive, suggesting the existence of a pre-C.1.2 strain that encode the common mutations shared by these two groups of subvariants. This hypothetical strain possesses many more mutations than its parental C.1 strain and might have emerged at the beginning of 2021. Among over 4 million genomes available from the GISAID sequence database, there are no apparent intermediate strains that bridge the large gap C.1 and pre-C.1.2 variants, suggesting accelerated evolution during emergence of the pre-C.1.2 variant.

**Figure 5.**
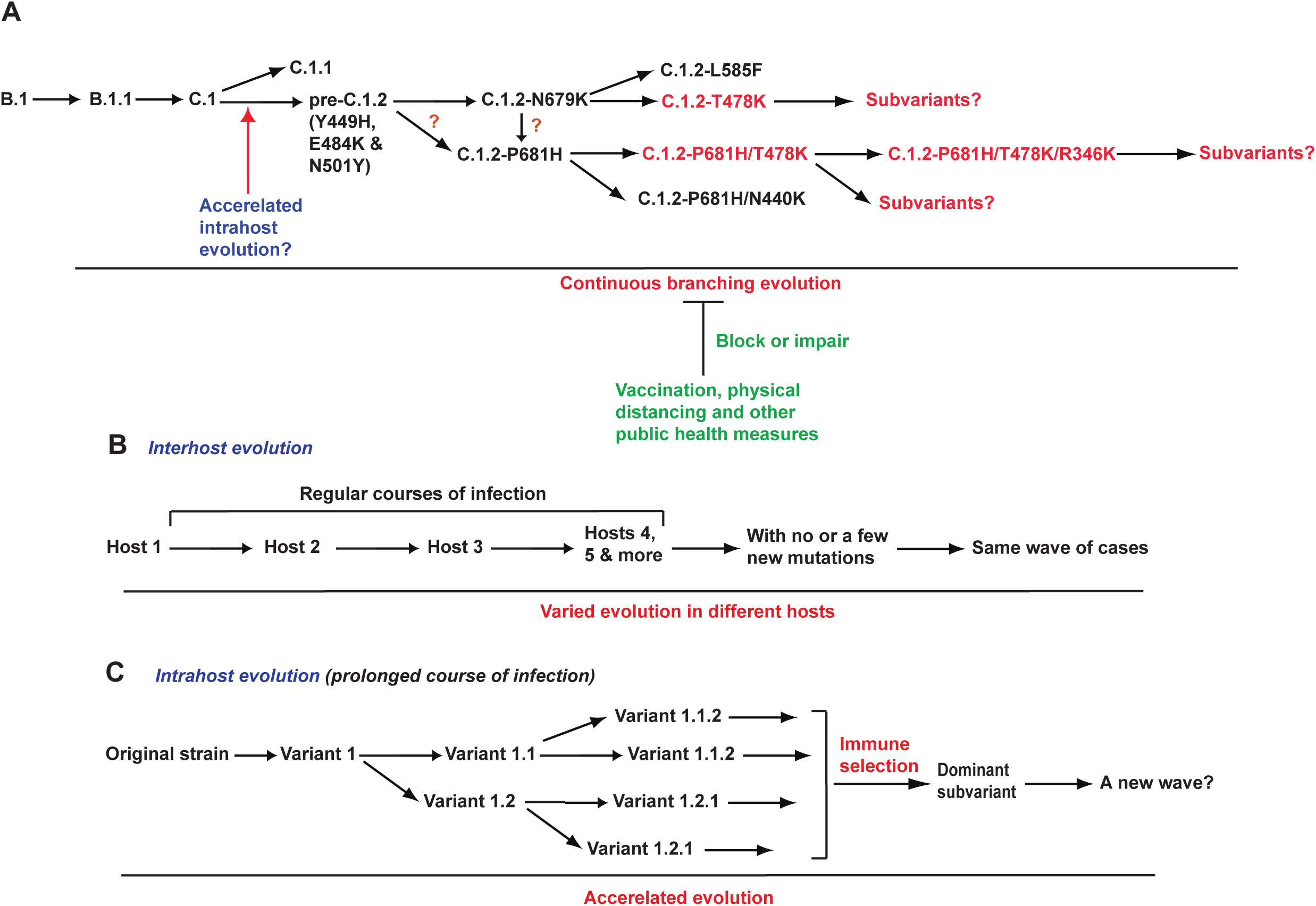
Schematic representation showing how SARS-COV-2 evolves continuously and generates different variants and subvariants. **(A)** Schematic representation showing how different variants evolve and gain additional substitutions. (**B-C**) Hypothetical models explaining two modes of evolution that drive the pandemic. In a majority of COVID-19 cases showing regular courses of infection (B), the evolution speed is rather limited, gaining no or a few mutations through interhost transmission. Such evolutionary mode would not change the course of the pandemic. Different from this is an accelerated mode of intrahost evolution (C). This mode explains sudden emergence of C1.2 variant and perhaps also alpha variant in September 2020. In a minority of COVID-19 cases showing prolonged courses of infection perhaps due to compromised immunity (C), virus evolution is accelerated and a dominant variant is selected, which then changes the course of the pandemic.

Hypothetical models in Fig. 5B-C explain two modes of evolution that drive the pandemic. In a majority of COVID-19 cases showing regular courses of infection (Fig. 5B), the evolution speed is rather limited, gaining no or a few mutations through interhost transmission. If such mutations do not gain substantial improvement in viral fitness, such evolutionary mode would not change the course of the pandemic. Examples are the initial phase of the pandemic and the waves of alpha variant cases. However, if some of these mutations confer much improved evolutionary advantage, such a mode of evolution may also change the course of the pandemic. One example is the acquisition of the spike substitution D614G and the NSP12 substitution P323L, yielding the parental strain for more than >99% of all COVID-19 cases. One important question is how C.1 suddenly gained so many mutations to produce the pre-C.1.2 strain. This might have involved an accelerated mode of intra-host evolution (Fig. 5C). This mode explains sudden emergence of alpha variant in September 2020 and perhaps also the other variants of concern. In a minority of COVID-19 cases showing prolonged courses of infection perhaps due to compromised immunity (Fig. 5C), virus evolution is accelerated and a dominant variant is selected. This variant is passed on to a different individual and additional mutations are gained to generate variants with potential to cause new waves of cases, which then changes the course of the pandemic.

There are genomes encoding different C.1.2 subvariants, with the first case identified in March 2021 (Fig. 3). The genomes possess an average of 46-47 mutations per genome. This mutation load is much higher than α [2], β [3] and γ (P.1) [12] variants of concern. The most mutated δ subvariants identified so far encode an average of 46-50 mutations per genome [13-16]. Thus, C.1.2 variant is one of the most mutated SARS-COV-2 lineages. Based on its genomes identified from June to September 2021, C.1.2 is more virulent than β variant but much weaker than δ variant. Related to this, Y449H reduces ACE2 receptor-binding [7]. Therefore, C.1.2 variant may not as virulent as its high mutation load suggests.

## Supporting information

Acknowledgement table on the GISAID genomes used in this study

## ACKNOWLEDGEMENT

I gratefully acknowledge the GISAID database for maintenance SARS-COV-2 viral genomes and the investigators for the valuable genomes sequences used in this work (see the supplementary section for details). I am grateful to Professor Federico M. Giorgi at University of Bologna, Italy for generosity to allow the access to the Coronapp server. This work was supported by funds from Canadian Institutes of Health Research (CIHR), Natural Sciences and Engineering Research Council of Canada (NSERC) and Compute Canada (to X.J.Y.).

## MATERIALS AND METHODS

### SARS-COV-2 virus sequence files, mutation profiling and phylogenetic analysis

Two substitutions (NSP3_Y428I and Spike_Y449H) were used simultaneously to search through over 3 million SARS-COV-2 genomes in the GISAID database on September 02, 2021. This search yielded 130 genomes, with 116 of them containing information on the age of the infected patients. The age information was downloaded for calculation of age distribution. Gender information is available for many of them and was used to calculate gender-specific association. CoVsurver (https://www.gisaid.org/epiflu-applications/covsurver-mutations-app/) was used to analyze mutations on the index case and some representative entries.

A Fasta file containing the 130 genomes was then downloaded from the GISAID database. During downloading, empty spaces in the fasta headers were replaced by underscores because such spaces make the files incompatible for subsequent sequence alignment and phylogenetic analysis on a Mac computer. To shorten the Fasta headers, the Find/Replace_All function of TextEdit (version 1.16) was used to delete “Cov19/” and “/2021” at the beginning and middle of the headers, respectively. In addition, “/” and “|” symbols in the headers are incompatible for sequence alignment and phylogenetic analysis, so they were also replaced with underscores via the Find/Replace_All function of TextEdit. The cleaned Fasta file was first used for mutational profiling via Coronapp (http://giorgilab.unibo.it/coronannotator/), a web-based mutation annotation application [10,11].

The cleaned Fasta file was then uploaded to SnapGene (version 5.3.2) for multisequence analysis via the MAFFT tool. To maximize coverage, all sequences were included in the analysis. After the alignment, the 5’ and 3’ were trimmed to make over 95% genomes possess the same length. The aligned sequence was exported as a Fasta file. Each header in this file contains “(0 bp)” some adjacent empty spaces, which are incompatible for subsequent phylogenetic analysis and were thus deleted via the Find/Replace All function of TextEdit. The file was named as Y449H_T428I_Sept_02_2021_130.fa, which was transferred to a folder for RAxML-NG version 0.9.0 [18] for analysis via the Terminal mode on the Mac computer. The file was further cleaned via the command line: raxml-ng --check --msa Y449H_T428I_Sept_02_2021_130.fa --model GTR+G. This generated a file (Y449H_T428I_Sept_02_2021_130.fa.raxml.reduced.phy) for phylogenetic analysis via the following command line: raxml-ng --msa Y449H_T428I_Sept_02_2021_130.fa.raxml.reduced.phy --model GTR+G –prefix Y449H_T428I_Sept_02_2021_130 --threads 2 --seed 2. The phylogenetic analysis generated 20 maximum likelihood trees and one bestTree. FigTree V1.4.4 was used to open the 21 tree files for manual inspection, analysis and annotation. To guide the inspection, many of the 130 genomes were analyzed in batches to display mutations via CoVsurver (https://www.gisaid.org/epiflu-applications/covsurver-mutations-app/). In addition, results from mutation profiling via Coronapp [10] also served as a guide in manual inspection of the 21 trees. Furthermore, the time of case identification was also considered when a potential root was chosen. With these factors all considered, an optimal tree was selected from the 21 trees for re-rooting via FigTree (https://github.com/rambaut/figtree/releases/tag/v1.4.4). The resulting tree was exported for image processing via Adobe Photoshop and subsequent presentation via Illustrator.

### PyMol structural modeling

The PyMol molecular graphics system (version 2.4.2, https://pymol.org/2/) from Schrödinger, Inc. was used for downloading structure files from the PDB database for further analysis and image export. The images were cropped via Adobe Photoshop and further presentation using Illustrator.

## SUPPLEMENTAL INFORMATION

This section includes one Supplementary Figure and one acknowledgement table for the 130 GISAID genomes used in this work.

## SUPPLEMENTAL FIGURE LEGENDS

**Figure S1.**
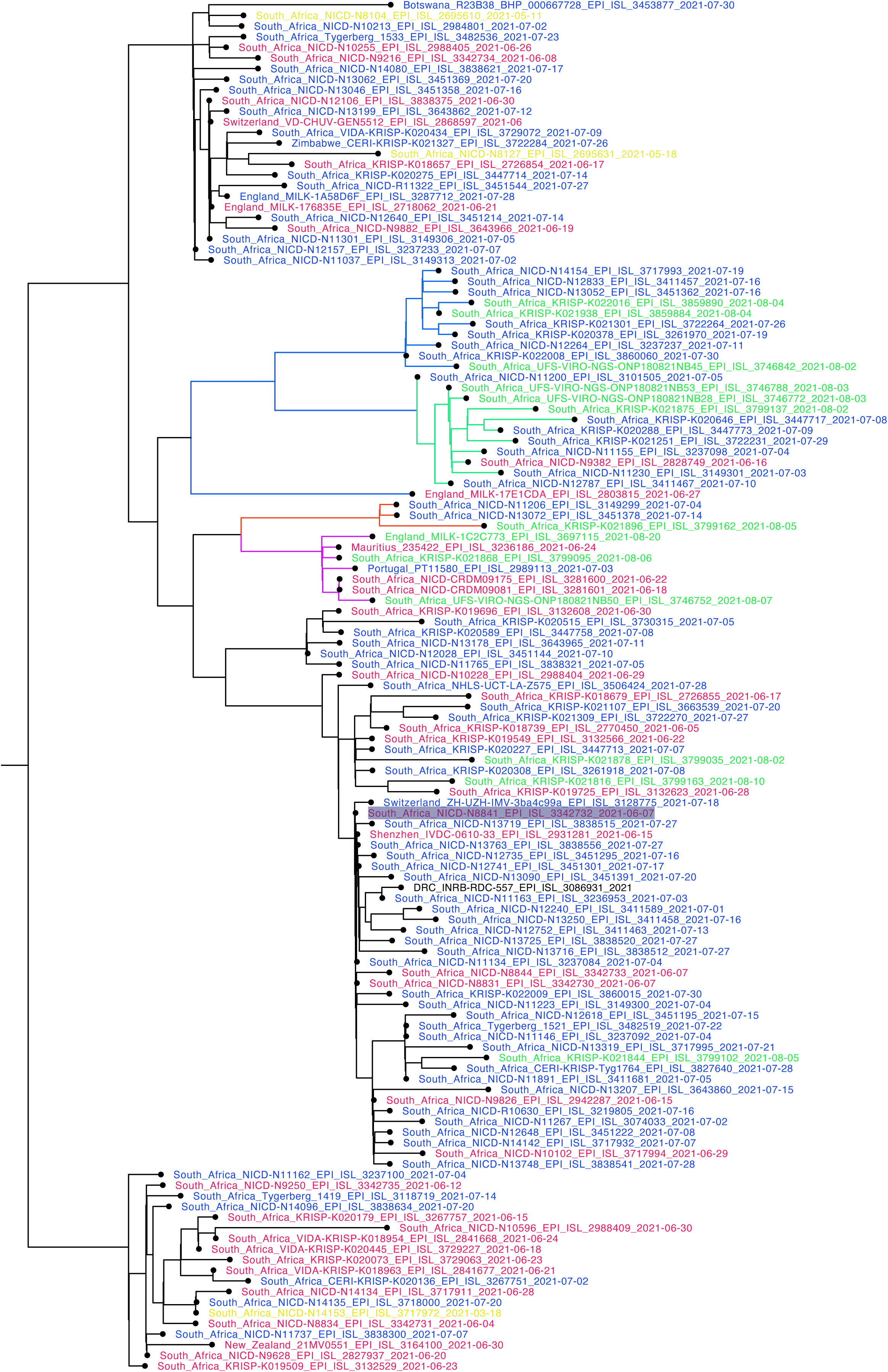
Phylogenetic analysis of 130 C.1.2 genomes from different countries around the world. A Fasta file containing the genomes was downloaded from the GISAID database and its Fast headers were modified via Textedt. The cleaned Fasta file was then uploaded to Snapgene for multisequence analysis via the MAFFT tool. The phylogenetic analysis package RAxML-NG was used to generate the maximum likelihood tree presented here, with strain names and GISAID accession numbers of the genomes shown. Genomes identified in different months are colored differently. The P681H and T478K branches are also colored. South_Africa_UFS-VIRO-NGS-ONP180821NB50_EPI_ISL_3746752_2021-08-07 and South_Africa_UFS-VIRO-NGS-ONP180821NB19_EPI_ISL_3746811_2021-08-04 are identical, so one of them were removed during the analysis by RAxML-NG. See Fig. 3 for comparison.

