## Supplementary material for "SARS-COV-2 C.1.2 variant is highly mutated but may possess reduced affinity for ACE2 receptor": Acknowledgement table on the GISAID genomes used in this study

We gratefully acknowledge the following Authors from the Originating laboratories responsible for obtaining the specimens, as well as the Submitting laboratories where the genome data were generated and shared via GISAID, on which this research is based.

All Submitters of data may be contacted directly via www.gisaid.org

Authors are sorted alphabetically.

| Accession ID | Originating Laboratory | Submitting Laboratory | Authors |
|---|---|---|---|
| EPI_ISL_3267757 | AHRI- Alex Sigal lab | KRISP, KZN Research Innovation and Sequencing Platform | Alex Sigal; Giandhari Jennifer; Mallory Bernstein; Naidoo Yeshnee; Pillay Sureshnee; San E. James; Sandile Cele; Tegally Houriiyah; Tshabuila Derek; Wilkinson Eduan; Yajna Ramphal; de Oliveira Tulio |
| EPI_ISL_3236186 | Airport Health Laboratory/Central Health Laboratory | Virology Department, Central Health Laboratory ,Victoria Hospital, Candos,Ministry of Health and Wellness, Mauritius | Bahadoor BS; Jannoo N; Manraj SS; Mathur H; Pattoo M; Ramuth M; Sonoo J; Sujeewon C |
| EPI_ISL_3827640 | Division of Medical Virology, National Health Laboratory Service (NHLS), Tygerberg Hospital / Stellenbosch University | CERI, Centre for Epidemic Response and Innovation, Stellenbosch University and CERI-KRISP, KZN Research Innovation and Sequencing Platform | Alvera Vorster; Bronwyn Kleinhans; Carel J van Heerden; Gert van Zyl; Giandhari Jennifer; Kamela Mahlakwane; Karabo Phadu; Mathilda Claassen; Naidoo Yeshnee; Pillay Sureshnee; Ren Veikondis; San James; Shannon Wilson; Susan Engelbrecht; Tania Stander; Tegally Houriiyah; Tongai Maponga; Tshiabuila Derek; Wilkinson Eduan; Wolfgang Preiser; Yajna Ramphal; de Oliveira Tulio |
| EPI_ISL_3118719, EPI_ISL_3482519, EPI_ISL_3482536 | Division of Medical Virology, National Health Laboratory Service (NHLS), Tygerberg Hospital / Stellenbosch University | Division of Medical Virology, National Health Laboratory Service (NHLS), Tygerberg Hospital / Stellenbosch University | Bronwyn Kleinhans; Gert van Zyl; Kamela Mahlakwane; Shannon Wilson; Susan Engelbrecht; Tania Stander; Tongai Maponga; Wolfgang Preiser |
| EPI_ISL_2868597 | EHNV | Laboratory of genomics and metagenomics | Claire Bertelli; Damien Jacot; Gilbert Greub; Sébastien Aeby; Trestan Pillonel |
| EPI_ISL_2942287, EPI_ISL_2988404, EPI_ISL_2988405, EPI_ISL_2988409 | HELEN JOSEPH LABORATORY | National Institute for Communicable Diseases of the National Health Laboratory Service | Amoako DG; Bhiman JN; Everatt J; Ismail A; Mahlangu B; Mnguni A; Mohale T; Ntuli N; Scheepers C |
| EPI_ISL_3342734, EPI_ISL_3342735 | Helen Joseph Laboratory | National Institute for Communicable Diseases of the National Health Laboratory Service | Amoako DG; Bhiman JN; Everatt J; Ismail A; Mahlangu B; Mnguni A; Mohale T; Ntuli N; Scheepers C |
| EPI_ISL_3451295, EPI_ISL_3451301, EPI_ISL_3838321, EPI_ISL_3838512, EPI_ISL_3838515, EPI_ISL_3838520, EPI_ISL_3838541, EPI_ISL_3838556 | | | |
| see above | KIMBERLEY LABORATORY | National Institute for Communicable Diseases of the National Health Laboratory Service | Amoako DG; Bhiman JN; Everatt J; Ismail A; Mahlangu B; Mnguni A; Mohale T; Ntuli N; Scheepers C |
| EPI_ISL_3101505, EPI_ISL_3149299, EPI_ISL_3149300, EPI_ISL_3149301 | KOPANONG HOSPITAL | National Institute for Communicable Diseases of the National Health Laboratory Service | Amoako DG; Bhiman JN; Everatt J; Ismail A; Mahlangu B; Mnguni A; Mohale T; Ntuli N; Scheepers C |
| EPI_ISL_3411463, EPI_ISL_3411467, EPI_ISL_3643966 | KOPANONG LABORATORY | National Institute for Communicable Diseases of the National Health Laboratory Service | Amoako DG; Bhiman JN; Everatt J; Ismail A; Mahlangu B; Mnguni A; Mohale T; Ntuli N; Scheepers C |
| EPI_ISL_3237237 | LANCET LABORATORY | National Institute for Communicable Diseases of the National Health Laboratory Service | Amoako DG; Bhiman JN; Everatt J; Ismail A; Mahlangu B; Mnguni A; Mohale T; Ntuli N; Scheepers C |
| EPI_ISL_3149313, EPI_ISL_3451358, EPI_ISL_3451362, EPI_ISL_3451369, EPI_ISL_3451378, EPI_ISL_3451391 | LEBOWAKGOMO LABORATORY | National Institute for Communicable Diseases of the National Health Laboratory Service | Amoako DG; Bhiman JN; Everatt J; Ismail A; Mahlangu B; Mnguni A; Mohale T; Ntuli N; Scheepers C |
| EPI_ISL_3506424 | Lancet | NHLS/UCT | Arash Iranzadeh; Bruna Galvao; Carolyn Williamson; Deelan Doolabh; Diana Hardie; Gert Marais; Innocent Mudau; Lynn Tyers; Marvin Hsiao; Rageema Joseph; Sisonke; Stephen Korsman |
| EPI_ISL_2718062, EPI_ISL_2803815, EPI_ISL_3287712, EPI_ISL_3697115 | Lighthouse Lab in Milton Keynes | Wellcome Sanger Institute for the COVID-19 Genomics UK (COG-UK) Consortium | Cordelia Langford; David K. Jackson; Dominic Kwiatkowski; Ewan Harrison; Ian Johnston; Jeffrey Barrett; John Sillitoe on behalf of the Wellcome Sanger Institute COVID-19 Surveillance Team; Roberto Amato; Sonia Goncalves; The Lighthouse Lab in Milton Keynes and Alex Alderton |
| EPI_ISL_2827937, EPI_ISL_3451144, EPI_ISL_3717995, EPI_ISL_3838300 | MANKWENG LABORATORY | National Institute for Communicable Diseases of the National Health Laboratory Service | Amoako DG; Bhiman JN; Everatt J; Ismail A; Mahlangu B; Mnguni A; Mohale T; Ntuli N; Scheepers C |
| EPI_ISL_3164100 | Middlemore Hospital | Institute of Environmental Science and Research (ESR) | Anja Werno; Antje van der Linden; Arlo Upton; Chris Mansell; David Hammer; Dragana Drinkovic; Erasmus Smit; Gary McAuliffe; Hana Sofia Andersson; Hermes Perez; James Ussher; Jill Sherwood; Jing Wang; Joep de Ligt; Josh Freeman; Julia Howard; Juliet Elvy; Lauren Jelly; Mary DeAlmeida; Matt Blakiston; Matt Storey; Matthew Rogers; Max Bloomfield; Michael Addidle; Michelle Balm; Muhammad Faisal; Nikki Freed; Olin Silander; Olivia Stroeven; Rachel Boyle; Sally Roberts; SallyAnn Harbison; Sarah Jefferies; Sharmini Muttaiyah; Susan Morpeth; Susan Taylor; Timothy Blackmore; Vani Sathyendran; Veronica Playle; Virginia Hope; Xiaoyun Ren |
| EPI_ISL_2726854, EPI_ISL_2726855, EPI_ISL_2770450 | NHLS Charlotte Maxeke Johannesburg Academic Hospital and the University of the Witwatersrand | KRISP, KZN Research Innovation and Sequencing Platform | Bulelani Manene; Florette Treurnicht; Giandhari Jennifer; Kathleen Subramoney; Naidoo Yeshnee; Pillay Sureshnee; San James; Tegally Houriiyah; Tshabuila Derek; Wilkinson Eduan; Yajna Ramphal; de Oliveira Tulio |
| EPI_ISL_3132529, EPI_ISL_3132566, EPI_ISL_3132623 | NHLS Charlotte Maxeke Johannesburg Academic Hospital and the University of the Witwatersrand | KRISP, KZn Research Innovation and Sequencing Platform | Bulelani Manene; Florette Treurnicht; Giandhari Jennifer; Kathleen Subramoney; Naidoo Yeshnee; Pillay Sureshnee; San James; Tegally Houriiyah; Tshabuila Derek; Wilkinson Eduan; Yajna Ramphal; de Oliveira Tulio |
| EPI_ISL_3729063, EPI_ISL_3859884, EPI_ISL_3859890, EPI_ISL_3860015, EPI_ISL_3860060 | NHLS Charlotte Maxeke Johannesburg Hospital | KRISP, KZN Research Innovation and Sequencing Platform | Emmanuel S; Florette Treurnicht; Giandhari J; Kathleen Subramoney; Naidoo Yeshnee; Pillay S; Tegally H; Tshabuila Derek; Wilkinson E; Yajna Ramphal; de Oliveira T |
| EPI_ISL_3722231, EPI_ISL_3722264, EPI_ISL_3722270, EPI_ISL_3799035, EPI_ISL_3799095, EPI_ISL_3799137, EPI_ISL_3799162 | | | |
| see above | NHLS Charlotte Maxeke Johannesburg Hospital | KRISP, KZn Research Innovation and Sequencing Platform | Emmanuel S; Florette Treurnicht; Giandhari J; Kathleen Subramoney; Naidoo Yeshnee; Pillay S; Tegally H; Tshabuila Derek; Wilkinson E; Yajna Ramphal; de Oliveira T |
| EPI_ISL_3746752, EPI_ISL_3746772, EPI_ISL_3746788, EPI_ISL_3746811, EPI_ISL_3746842 | NHLS Universitas Academic | UFS Virology | D Goedhals; Emmanuel Ogunbayo; MM Nyaga; MT Mogotsi; P Nthiga; PA Bester; T de Oliveira |
| EPI_ISL_3447717, EPI_ISL_3447758 | NHLS_IALCH | KRISP, KZN Research Innovation and Sequencing Platform | Giandhari Jennifer; Naidoo Yeshnee; Pillay Sureshnee; San James; Tegally Houriiyah; Tshiabuila Derek; Wilkinson Eduan; Yajna Ramphal; de Oliveira Tulio |
| EPI_ISL_3447713, EPI_ISL_3447714, EPI_ISL_3447773 | NHLS_JHB | KRISP, KZN Research Innovation and Sequencing Platform | Bulelani Manene; Florette Treurnicht; Giandhari Jennifer; Kathleen Subramoney; Naidoo Yeshnee; Pillay Sureshnee; San James; Tegally Houriiyah; Tshiabuila Derek; Wilkinson Eduan; Yajna Ramphal; de Oliveira Tulio |
| EPI_ISL_3261918 | NHLS_JHB (Ship 4) | KRISP, KZN Research Innovation and Sequencing Platform | Giandhari Jennifer; Naidoo Yeshnee; Pillay Sureshnee; San James; Tegally Houriiyah; Tshabuila Derek; Wilkinson Eduan; Yajna Ramphal; de Oliveira Tulio |
| EPI_ISL_3261970 | NHLS_VIRO | KRISP, KZN Research Innovation and Sequencing Platform | Giandhari Jennifer; Naidoo Yeshnee; Pillay Sureshnee; San James; Tegally Houriiyah; Tshabuila Derek; Wilkinson Eduan; Yajna Ramphal; de Oliveira Tulio |
| EPI_ISL_3663539 | National Health Laboratory Service, South Africa | KRISP, KZN Research Innovation and Sequencing Platform | Giandhari Jennifer; Naidoo Yeshnee; Pillay Sureshnee; San James; Tegally Houriiyah; Tshabuila Derek; Wilkinson Eduan; Yajna Ramphal; de Oliveira Tulio |
| EPI_ISL_3132608, EPI_ISL_3730315, EPI_ISL_3799102, EPI_ISL_3799163 | National Health Laboratory Service, South Africa | KRISP, KZn Research Innovation and Sequencing Platform | Emmanuel S; Giandhari J; Giandhari Jennifer; Naidoo Yeshnee; Pillay S; Pillay Sureshnee; San James; Tegally H; Tegally Houriiyah; Tshabuila Derek; Tshiabuila Derek; Wilkinson E; Wilkinson Eduan; Yajna Ramphal; de Oliveira T; de Oliveira Tulio |
| EPI_ISL_2984801, EPI_ISL_3149306, EPI_ISL_3219805, EPI_ISL_3281600, EPI_ISL_3281601, EPI_ISL_3451544, EPI_ISL_3717994 | | | |
| see above | National Institute for Communicable Diseases of the National Health Laboratory Service | National Institute for Communicable Diseases of the National Health Laboratory Service | Amoako DG; Bhiman JN; Everatt J; Ismail A; Mahlangu B; Mnguni A; Mohale T; Ntuli N; Scheepers C; Sisonke Team |
| EPI_ISL_3722284 | National Microbiology Reference Laboratory, Ministry of Health, Harare, Zimbabwe | CERI, Centre for Epidemic Response and Innvoation, Stellenbosch University and KRISP, KZN Research Innovation and Sequencing Platform, UKZN. | Agnes Juru; Air Comodor Dr J. Chimedza; Charles Nyagupe; Dr Raiva Simbi; Emmanuel SJ; Giandhari J; Hlanai Gumbo; Kenneth Maeka; Naidoo Y; Pillay S; Tapfumanei Mashe; Tatenda Takawira; Tegally H; Wilkinson E; de Oliveira T |
| EPI_ISL_3411457, EPI_ISL_3411458, EPI_ISL_3451222 | POLOKWANE LABORATORY | National Institute for Communicable Diseases of the National Health Laboratory Service | Amoako DG; Bhiman JN; Everatt J; Ismail A; Mahlangu B; Mnguni A; Mohale T; Ntuli N; Scheepers C |
| EPI_ISL_3838375 | PORT ELIZABETH LABORATORY | National Institute for Communicable Diseases of the National Health Laboratory Service | Amoako DG; Bhiman JN; Everatt J; Ismail A; Mahlangu B; Mnguni A; Mohale T; Ntuli N; Scheepers C |
| EPI_ISL_3453877 | Palapye Primary Hospital Laboratory | Botswana Harvard HIV Reference Laboratory | Boitumelo J.L Zuze; Botshelo Radibe; Dorcas Maruapula; Joseph Makhema; Keoratile Ntshambiwa; Legodile Kooepile; Letsibogo Gaoraelwe; Madisa Mine; Modisa Motswaledi; Mosepele Mosepele; Ontlametse T. Bareng; Roger Shapiro; Shahin Lockman; Sikhulile Moyo; Simani Gaseitsiwe; Thela Tefelo; Thongbotho Mphoyakgosi; Wonderful T. Choga |
| EPI_ISL_3717911, EPI_ISL_3717932, EPI_ISL_3717972, EPI_ISL_3717993, EPI_ISL_3718000 | Pathcare | National Institute for Communicable Diseases of the National Health Laboratory Service | Amoako DG; Bhiman JN; Everatt J; Ismail A; Mahlangu B; Mnguni A; Mohale T; Ntuli N; Scheepers C; Sisonke Team |
| EPI_ISL_2828749, EPI_ISL_3411681 | Pathcare Vaal | National Institute for Communicable Diseases of the National Health Laboratory Service | Amoako DG; Bhiman JN; Everatt J; Ismail A; Mahlangu B; Mnguni A; Mohale T; Ntuli N; Scheepers C |
| EPI_ISL_3838621, EPI_ISL_3838634 | Private/Pathcare | National Institute for Communicable Diseases of the National Health Laboratory Service | Amoako DG; Bhiman JN; Everatt J; Ismail A; Mahlangu B; Mnguni A; Mohale T; Ntuli N; Scheepers C; Sisonke Team |

| | | | |
|---|---|---|---|
| EPI_ISL_2695610, EPI_ISL_2695631, EPI_ISL_3237233, EPI_ISL_3643860, EPI_ISL_3643862, EPI_ISL_3643965 | ROB FERREIRA LABORATORY | National Institute for Communicable Diseases of the National Health Laboratory Service | Amoako DG; Bhiman JN; Everatt J; Ismail A; Mahlangu B; Mnguni A; Mohale T; Ntuli N; Scheepers C |
| EPI_ISL_2989113 | SESARAM | Instituto Nacional de Saude (INSA) | Borges et al |
| EPI_ISL_2931281 | Shenzhen Center for Disease Control and Prevention | National Institute for Viral Disease Control and Prevention, China CDC | Can Zhu; Kai Nie; Long Chen; Peihua Niu; Renli Zhang; Shaoyu Deng and Yaqing He; Weihua Wu; Xinyi Wei; Yue Li |
| EPI_ISL_3267751 | Sisonke - BARC SA | CERI, Centre for Epidemic Response and Innovation, Stellenbosch Univeristy & KRISP, KZN Research Innovation and Sequencing Platform | Giandhari Jennifer; Naidoo Yeshnee; Pillay Sureshnee; San James; Tegally Houriiyah; Tshabuila Derek; Wilkinson Eduan; Yajna Ramphal; de Oliveira Tulio |
| EPI_ISL_3128775 | Stadtspital Triemli | Institute of Medical Virology | Alexandra Trkola; Annette Audigé; Catharine Aquino; Cyril Shah; Daniel Ehrsam; Gabriela Ziltener; Guido Bloemberg; Hubert Rehrauer; Isabel Stürmer; Joel Wirz; Jon Huder; Jürg Böni; Kevin Steiner; Maria Grünberg; Maryam Zaheri; Michael Huber; Riccarda Capaul; Stefan Schmutz; Verena Kufner; Weihong Qi |
| EPI_ISL_3074033, EPI_ISL_3236953, EPI_ISL_3237084, EPI_ISL_3237092, EPI_ISL_3237098, EPI_ISL_3237100, EPI_ISL_3451195, EPI_ISL_3451214 | | | |
| see above | TAMBO MEMORIAL LABORATORY | National Institute for Communicable Diseases of the National Health Laboratory Service | Amoako DG; Bhiman JN; Everatt J; Ismail A; Mahlangu B; Mnguni A; Mohale T; Ntuli N; Scheepers C |
| EPI_ISL_3342730, EPI_ISL_3342731, EPI_ISL_3342732, EPI_ISL_3342733 | Tambo Memorial | National Institute for Communicable Diseases of the National Health Laboratory Service | Amoako DG; Bhiman JN; Everatt J; Ismail A; Mahlangu B; Mnguni A; Mohale T; Ntuli N; Scheepers C |
| EPI_ISL_3411589 | University Pretoria | National Institute for Communicable Diseases of the National Health Laboratory Service | Amoako DG; Bhiman JN; Everatt J; Ismail A; Mahlangu B; Mnguni A; Mohale T; Ntuli N; Scheepers C |
| EPI_ISL_3729072, EPI_ISL_3729227 | Vaccines and Infectious Diseases Analytics Research Unit (VIDA) | KRISP, KZN Research Innovation and Sequencing Platform | Baillie Vicky; Giandhari Jennifer; Madhi Shabir; Naidoo Yeshnee; Pillay Sureshnee; San James; Tegally Houriiyah; Tshabuila Derek; Wilkinson Eduan; Yajna Ramphal; de Oliveira Tulio; du Plessis Jeanine |
| EPI_ISL_3086931 | Viral Respiratory Lab, National Institute for Biomedical Research (INRB) | Pathogen Sequencing Lab, National Institute for Biomedical Research (INRB) | Allison Black; Amuri Aziza; Andrew Rambaut; Catherine Pratt; Eddy Kinganda-Lusamaki; Edith Nkwembe; Emmanuel Lokilo Lofiko; Francisca Muyembe Mawete; Gabriel Kabamba; Ian Goodfellow; James Hadfield; Jean Claude Makangara; Jean-Jacques Muyembe Tamfum; Josh Quick; Kristian Andersen; Matthias Pauthner; Michael Wiley; Nick Loman; Placide Mbala-Kingebeni; Raphaël Lumembe; Steve Ahuka-Mundeke; Trevor Bedford |
| EPI_ISL_2841668, EPI_ISL_2841677 | Wits Vaccines & Infectious Diseases Analytics (VIDA) Research Unit | KRISP, KZn Research Innovation and Sequencing Platform | Baillie Vicky; Giandhari Jennifer; Madhi Shabir; Naidoo Yeshnee; Pillay Sureshnee; San James; Tegally Houriiyah; Wilkinson Eduan; de Oliveira Tulio; du Plessis Jeanine |
